# The yeast GRASP Grh1 displays a high polypeptide backbone mobility along with an amyloidogenic behavior

**DOI:** 10.1101/254144

**Authors:** N. A. Fontana, R. Fonseca-Maldonado, L.F.S Mendes, L. P. Meleiro, A. J. Costa-Filho

**Author notes:** Corresponding author: Antonio J. Costa-Filho, Av. Bandeirantes 3900, 14040-901, Ribeirão Preto, SP, Brazil. These authors contributed equally to this work (N. A. Fontana, R. Fonseca-Maldonado).

## Abstract

GRASPs are proteins involved in cell processes that seem paradoxical, such as being responsible for shaping the Golgi cisternae and also involved in unconventional secretion mechanisms that bypass the Golgi, among other functions in the cell. Despite its involvement in several relevant cell processes, there is still a considerable lack of studies on full-length GRASPs. Our group has previously reported an unexpected behavior of the full-length GRASP from the fungus *C. neoformans*: its intrinsically-disordered characteristic. Here, we generalize this finding by showing that is also observed in the GRASP from the yeast *S. cerevisae* (Grh1), which strongly suggests it may be a general property within the GRASP family. Furthermore, Grh1 is also able to form amyloid fibrils either upon heating or when submitted to changes in the dielectric constant of its surroundings, a condition that is experienced by the protein when in close contact with membranes of cell compartments, such as the Golgi apparatus. Intrinsic disorder and amyloid fibril formation can thus be two structural properties exploited by GRASP during its functional cycle.

## 1. Introduction

The Golgi complex is composed of a series of cisternal membranes opposed to one another to form stacks (Rambourg et al., 1981). In mammalian cells, the stacks are linked at their edges by tubules to form a ribbon-like structure (Barr et al., 1997; Rambourg and Clermont, 1990). An assay that blocks cisternal stacking in postmitotic events was the basis for the discovery of the two proteins known as Golgi Reassembly and Stacking Proteins (GRASP65 and GRASP55) (Barr et al., 1997; Shorter et al., 1999). Furthermore, other functions of GRASPs have already been pointed out, such as chaperoning and transport of some proteins, participation in cell apoptosis, Golgi reorientation during cell migration, unconventional protein secretion, and, during mitosis, as a possible G2/M checkpoint (Vinke et al., 2011).

GRASP structure is divided into two regions: an N-terminal half, called GRASP domain, which contains two putative PDZ domains (Kinseth et al., 2007) and the second half (the C-terminal region), rich in proline, serine, glutamine and asparagine residues, also known as the SPR domain (Feng et al., 2013; Truschel et al., 2011; Wang et al., 2005). The formation of the Golgi ribbon-like structure requires membrane bridging by the dimeric state of the GRASP domain (Feng et al., 2013; Truschel et al., 2011). In mammalian and Drosophila, GRASPs are tightly associated with the Golgi membranes via an N-myristoylation of the residue Gly_2_ (Barr et al., 1997; Kondylis et al., 2005) and, in yeasts, via an acetylated amphipathic helix (Behnia et al., 2007). The association of GRASP65 also depends on a Golgi receptor, identified as the coiled-coil protein called GM130 (Barr et al., 1998). The dual membrane association is important for the correct *trans* dimerization, a necessary step in the stack formation (Bachert et al., 2010; Heinrich et al., 2014).

Details of the involvement of GRASPs in membrane trafficking and other functions in mammalian cells have been reported by researchers using model organisms, such as the yeast *Saccharomyces cerevisiae*. Although *Saccharomyces cerevisiae* has the basic organization of its Golgi cisternae, only 40% of the cisternae are in stacks and the stacks are never found linked to each other (Vinke et al, 2011). This budding yeast contains a single GRASP homolog, known as Grh1, which localizes in compartments of the early secretory pathway (Levi et al., 2010). Grh1 is analog to GRASP65 and forms a complex with a coiled-coil protein, Bug-1, that shares structural features with GM130. The complex Grh1-Bug1 is involved in membrane trafficking, contributes to the formation of the cis-Golgi (Behnia et al., 2007) and, although dispensable for conventional secretion, is essential for the unconventional secretion of ACBP1 (Giuliani et al., 2011). Furthermore, Grh1 interacts with the dimer formed by Sec23 and Sec24, protein components of the COPII coat, an event necessary for the fusion of vesicles derived from ER with Golgi membranes (Behnia et al., 2007).

Here, we present the first structural characterization of the yeast GRASP Grh1. We investigated the biophysical and biochemical features of Grh1 and the GRASP domain only (called here DGRASP) by circular dichroism (CD), fluorescence and optic spectroscopies, differential scanning calorimetry (DSC), computational predictions and established that Grh1 is a molten globule-like protein, making it a member of the collapsed intrinsically disordered protein (IDP) family. IDPs are proteins involved in a large set of functions and characterized by regions of high polypeptide mobility, and without a well-defined 3D structure (Dunker et al., 2001; Dunker et al., 2008). These proteins have been grouped into two broad structural classes: (1) collapsed (molten globule-like) and (2) extended (coil-like and pre-molten globule-like) (Permyakov et al., 2008). The structural flexibility of IDPs allows a broad functional repertoire and a number of interaction partners (Uversky, 2009) to act and to influence protein function in different processes, such as transcriptional regulation, translation, cellular signal transduction, and storage of small molecules (Perticaroli et al., 2014).

Alongside with its disorder, Grh1 also shows an unexpected feature. We report here our findings on the amyloidogenic behavior of this GRASP. They are derived from CD, fluorescence using a specific dye, and Congo Red absorbance experiments. The results obtained from this wide range of techniques led us to the conclusion that Grh1 can form amyloid-like structures in conditions that could be reasonably found in the cell. More than that, we showed that the DGRASP, which is the most conserved region along GRASP family, is sufficient for the fiber formation. Our results suggest that this could be a general feature of GRASPs.

## 2. Material and Methods

### 2.1. Bioinformatics Tools

The aggregation prediction was done in the AGGRESCAN server (Conchillo-Solé et al., 2007), using a 5-residue window. The disorder prediction was done using the DisEMBL (Iakoucheva; Dunker, 2003) and the PONDR-FIT (Xue et al., 2010) servers.

### 2.2. Protein expression and purification

Genomic DNA of a strain of *Saccharomyces cerevisiae* was used as the template for PCR amplification of the gene encoding Grh1 (Gene ID: 852129) using primers Grh1F (5’-CCCGGATCCTTTAGAATAGCTAAAAACCTCGTACGG-3’) and Grh1R (5’-GGGTTCGAATTAATCAGAGGATGACTGTTTTTGTGGT‐ 3’). The PCR reaction was carried at 94 ºC for 3 min, followed by 30 cycles of 94 ºC for 1 min, 50 ºC for 1 min, 72 ºC for 1 min and final incubation at 72 °C for 10 min. The PCR product was digested with BamHI and HindII (recognition sites underlined in the oligonucleotide sequences) and cloned into the plasmid pETSUMO. The resulting construct (pETSUMO-Grh1) was transformed into DH5α *Escherichia coli*, and the plasmid DNA was purified and sequenced. *E. coli* Rosetta (DE3) cells (Novagen, Darmstadt, Germany) transformed with pETSUMO-Grh1 were grown at 37°C and 200 rpm agitation until reaching an OD 600 nm of 0.6 in 2 L shake flasks containing 1 L LB medium supplemented with 40 µg.mL^−1^ kanamycin and 34 µg.mL^−1^ chloramphenicol. The expression was carried out for 21 h and induced with 0.5 mM IPTG at 18 °C and 200 rpm agitation. The cells were harvested and transferred to 20 mL of lysis buffer (40 mmol.L^-1^ HEPES pH 8.0, 300 mmol.L^-1^ NaCl, and 10% Glycerol). After disruption by sonication, cell debris were removed by centrifugation, and the supernatant was applied to a nickel affinity column (Promega – Madison, USA). The column was washed with buffer containing 40 mmol.L^-1^ HEPES pH 8.0, 300 mmol.L^-1^ NaCl, 10% Glycerol supplemented with 25 mmol.L^-1^ imidazole and was eluted in the same buffer with 300 mmol.L^-1^ imidazole. The imidazole was removed by extensive washing using centrifugation in a Vivaspin column (GE Healthcare, Buckinghamshire, United Kingdom) and the sample was incubated for 3 h with recombinant ULP-1 protease followed by incubation in a nickel affinity chromatographic column to remove the SUMO protein and ULP-1 protease. The remaining contaminants were removed by size exclusion chromatography onto a Superdex 200 10/300 GL gel filtration column (GE Healthcare, Buckinghamshire, United Kingdom) in 40 mmol.L^-1^ HEPES pH 8.0, 300 mmol.L^-1^ NaCl, 10% Glycerol buffer. The purification of the GRASP domain (DGRASP) followed the same protocol, using a different reverse primer, with a stop codon at the end of the GRASP domain to exclude the SPR domain.

### 2.3. Circular Dichroism (CD)

Far-UV (190-260 nm) CD experiments were carried out in a Jasco J-815 CD Spectrometer (JASCO Corporation, Japan) equipped with a Peltier temperature control and using a quartz cell with a path length of 1 mm. Grh1 was in 10 mM sodium phosphate buffer, pH 8.0 and at final concentration of 5 µM. All far-UV CD spectra were recorded with a scan speed of 50 nm/min and at time response of 1 s. Chemical stability experiments were performed in the same buffer and increasing urea concentration (0-8.0 M). To investigate the effects of solvents Grh1 was incubated in aqueous methanol (MeOH) and acetonitrile (ACN) over a range of 0–50% solvent. The spectra were averaged, baseline-corrected and smoothed with a Savitsky-Golay filter using CDTools software (Lees et al., 2004). The processed spectra were deconvoluted by using the software Continll (Sreerama and Woody, 2000) with database 7 (van Stokkum et al., 1990) available in the DichroWeb analysis server (Whitmore and Wallace, 2004). The normalized root-mean-square deviation (NRMSD) goodness-of-fit parameter was always less than 0.15, suggesting that the calculated spectra are in agreement with the experimental data (Whitmore and Wallace, 2008).

### 2.4. Steady-State Fluorescence Spectroscopy

Intrinsic and extrinsic fluorescence were monitored using a Hitachi F-7000 fluorimeter equipped with a 150 W xenon arc lamp. The excitation and emission monochromators were set at 2.5 nm slit width in all experiments. The protein concentration was 5 µM for Grh1 and 7 µM for DGRASP in 40 mM Hepes, 150 mM NaCl, 10% glycerol. For tryptophan fluorescence experiments, the selective tryptophan excitation wavelength was set at 295 nm and the emission spectrum was monitored from 300 up to 400 nm. The fluorescence of tryptophan across chemical denaturation was measured in increasing concentrations of urea (0-7.5 M). For the ThT experiments, 15 µM of the dye solution was used along with the protein, excited at 440 nm. For the ANS experiments, a 250 µM solution was used, with excitation at 355 nm.

### 2.5. Congo Red Assay

The absorbance spectrum of Congo Red (CR) was monitored in the presence and in the absence of the protein, between 400 and 700 nm, with a Beckman DU 640 Uv-Vis Spectrometer. The CR was in a buffer solution as reported elsewhere (Nilson, 2004).

### 2.6. Transmission Electron Microscopy

The TEM experiments were performed in a JEOL JEM 2100. A 200 kV of acceleration voltage was applied in heated Grh1 samples (15 minutes at 45º C -150 ug/mL), deposited in a 15 mA discharged copper grid and stained with 2% uranyl acetate. The experiments were performed at the Brazilian Nanotechnology National Laboratory (LNNano), in Campinas-SP.

## 3. Results and discussions

### 3.1. Sequence and structure prediction

Grh1 is composed of 372 amino acids and the analysis of the protein family database using the Pfam program (Finn et al., 2014) predicted that the GRASP domain, including the two PDZ subdomains, comprises residues 1 to 280, and the SPR domain extends from residue 281 to 372. In addition, the sequence-based prediction of disordered regions (Figure 1) showed that the C-terminal domain and the central region of the PDZ subdomains (spanning 55% of the protein sequence) have high probability of being intrinsically disordered, a tendency already observed for the SPR and the PDZ subdomains within the GRASP family (Mendes et al, 2016).

**Figure 1:**
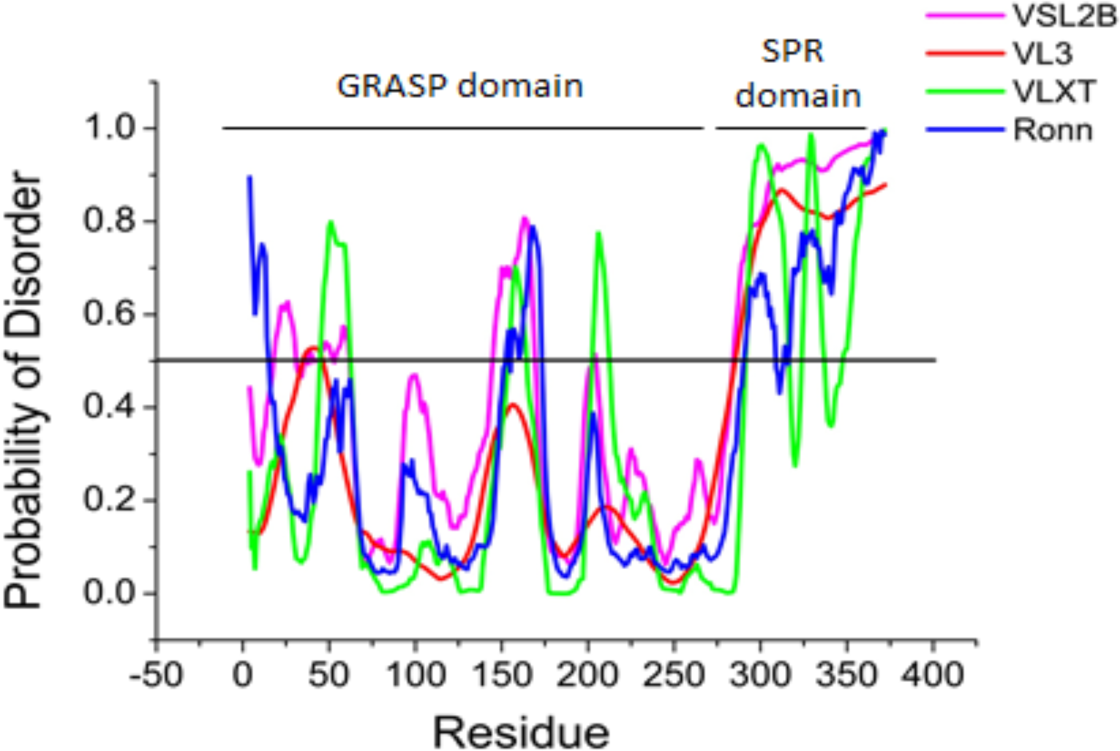
Predictions of intrinsically disordered regions in the Grh1 sequence using VSL2B (Magenta), VL3 (Red) VLXT, (Green) and Ronn (Blue). The black line indicates the threshold to be considered as a disordered region.

### 3.2. Structural Behavior in Solution

Unlike GRASP55 and GRASP65 (Feng et al., 2013; Truschel et al., 2011), full-length Grh1 was successfully expressed as a soluble, monodisperse protein in *E. coli* (Figure 2A). The theoretical molecular mass of the recombinant Grh1 is 41,119 Da, but SDS-PAGE analysis (Figure 2B) resulted in an apparent molecular mass of ca. 45,000 Da. This suggests that the amount of hydrophobic aminoacids that compose Grh1 is smaller than expected for well-structured proteins, a phenomenon similar to what was previously observed for other IDPs (Theillet et al., 2013). Size exclusion chromatography of the soluble protein on Superdex-200 column, whose result is shown in Figure 2B, indicates an apparent molecular mass of 45,200 Da. The differences between the expected molecular mass of Grh1 and the values determined from hydrodynamic methods is likely a consequence of the not-fully globular conformation of Grh1 in solution, which has been observed for other proteins rich in disordered regions (Chakrabortee et al., 2012), including the GRASP homologue in *C. neoformans* (Mendes et al, 2016). The chromatogram for the GRASP domain (DGRASP) is also presented in Figure 2B. We can see that it is eluted slightly after the full-length Grh1, which is expected since DGRASP lacks the SPR domain. Based on an elution curve calibrated with molecular mass standards (Suppl. Material, Figure S1), we conclude that Grh1 and its GRASP domain behave predominantly as a monomer in solution. This is different from the observed dimers in mammalian and rat GRASPs, which may be due to the lack, in the Grh1 primary sequence, of the residues involved in dimerization (Feng et al., 2013) and trans-oligomerization (Vinke et al., 2011) of GRASPs in mammalian and rat.

**Figure 2:**
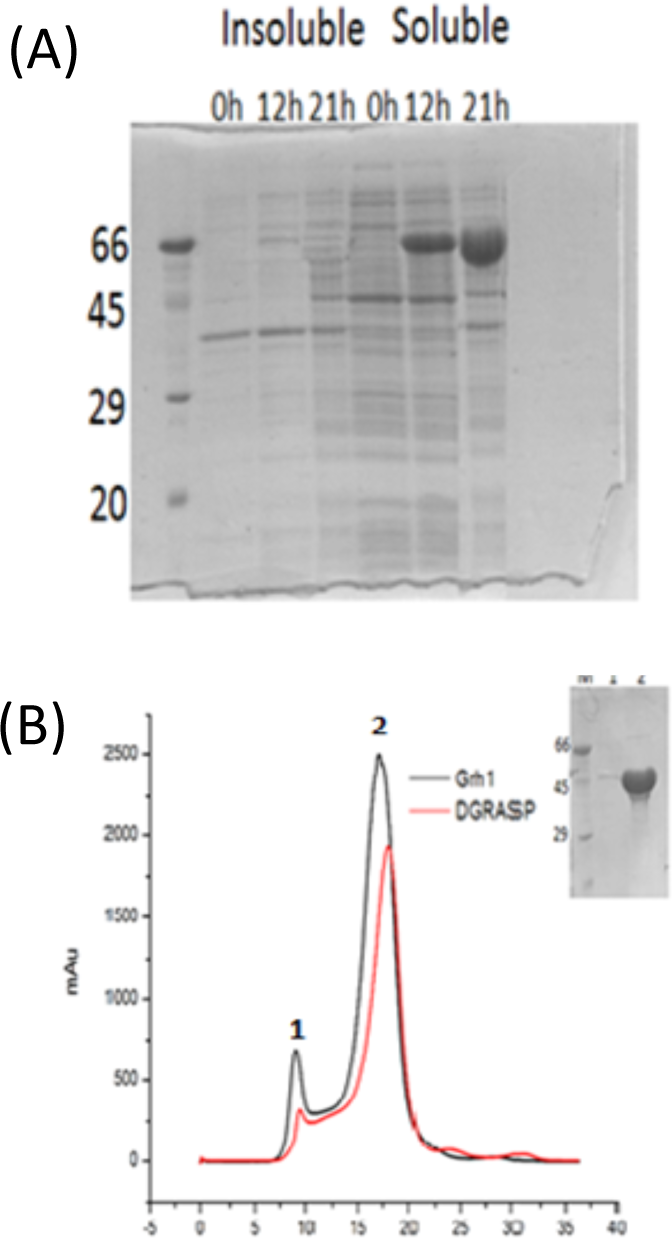
(A) SDS-PAGE monitoring the time course of Grh1 recombinant expression. Insoluble and soluble samples at specific times (0h, 12h 21h) after IPTG induction. (B) Size exclusion chromatography of Grh1 and DGRASP. The first peak represents aggregates of at least 45 molecules of Grh1 and the second peak, the elution of the monomeric Grh1 and DGRASP. Inset: SDS-PAGE of peaks 1 and 2. The pattern for the GRASP domain construct is the same observed for Grh1 (data not shown here).

The CD spectrum of Grh1 in aqueous solution (Figure 3) has a minimum around 204 nm and a poorly resolved and lower intensity peak at 222 nm, which are features typical of CD spectra of proteins with a high content of unordered structures (Uversky et al., 2009). However, the negative peak at 222 nm is an indication of some ordered elements. Although the intensity ratio of the peaks at 222 nm ([-4,264 deg.cm^2^.dmol^−1^) and 200 nm (−6,904 deg.cm^2^.dmol^−1^) in the CD spectrum of Grh1 is similar to values observed for other proteins in the pre-moltenglobule-like state, according to the “double wavelength” plot, [θ]_222_ vs. [θ]_200_ (Uversky, 2009), Grh1 does not fit perfectly as a natively unfolded protein based on the estimation of its secondary structure content (11.5% α-helix, 22.1% β-sheet, 17.4% turns, and 49.8% random coil). Comparing these results with those from the GRASP domain only, we observe that here the spectrum also presents a minimum around 200 nm and a low (even lower than for the whole protein) intensity peak at 222 nm, which suggests decreased ordering of the protein structure (Figure 3). In fact, when we subtract the DGRASP spectrum from that of Grh1, we have a spectrum that resembles that of a Poly(Pro)II conformation (Woody, 2009), which is expected based on the high content of prolines in the SPR domain.

**Figure 3:**
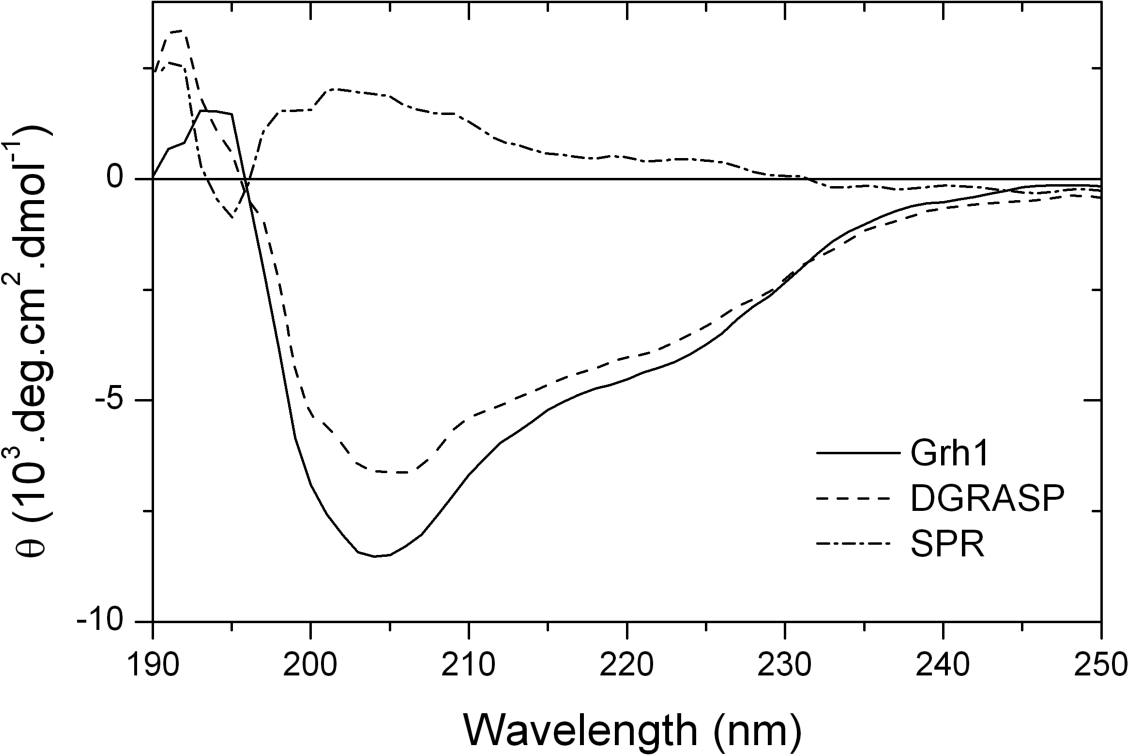
Far UV CD spectra of Grh1 (solid line), DGRASP (dotted line) and the SPR domain (dash line – Grh1 subtracted of DGRASP).

#### 3.2.1. Effects of Strong Denaturants

The urea-induced unfoldings of Grh1 and DGRASP were analyzed by CD and fluorescence spectroscopies. The unfolding monitored by CD spectroscopy (Figure 4) is a low cooperative transition as seen in the gradual change of the denatured fraction of the protein (f_d_) calculated from the molar ellipticity at 222 nm. The sigmoid-like transition is not as abrupt as expected for well-structured proteins of similar size (Uversky, 2009). The low steepness of the transition curve is typical of native molten globules or native coiled proteins and is due to the low percentage of secondary structure (Uversky, 2009; Mendes et al., 2016). We also obtain a low cooperative unfolding pattern for DGRASP, suggesting that the pattern observed for Grh1 does not come only from contributions of the SPR domain, but also from the GRASP domain (see below).

**Figure 4:**
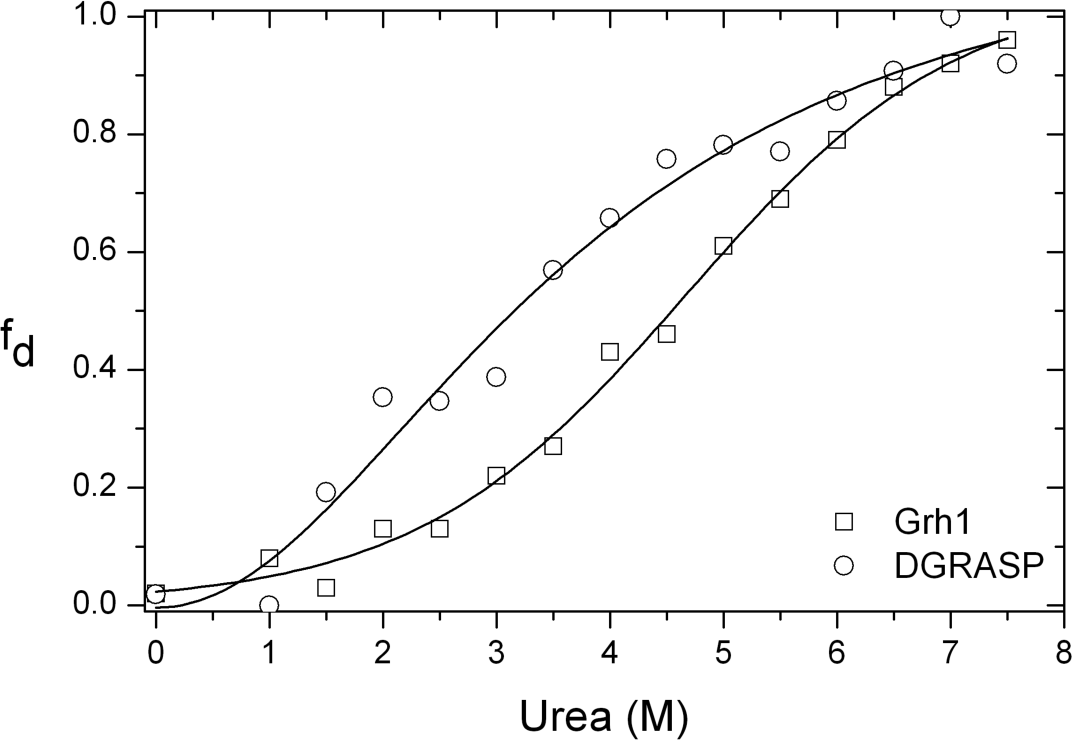
The unfolding fraction (fd) of Grh1 and DGRASP obtained from the CD intensity at 222 nm upon increasing concentrations of the denaturant. The solid lines are fits of the experimental data to a Boltzmann model.

The urea-induced unfolding was also monitored by using the wavelength of maximum fluorescence emission (λ_max_) of the tryptophan residues. Tryptophan in the aqueous environment has its maximum fluorescence emission around 350 nm, which is shifted to 320 nm when the aminoacid is placed in the hydrophobic core of proteins (Kazakov et al., 2009). For Grh1 in solution, the λ_max_ is centered at 344 nm indicating the tryptophan residues are exposed to the solvent. The fluorescence signal shows a red shift, in a cooperative transition, from 344 to 352 nm upon increasing urea concentrations (Figure 5A), indicating further exposure of the tryptophan residues and complete loss of the protein structure. Furthermore, at low concentrations of urea, the fluorescence anisotropy values remain unchanged up to a concentration of 2.5 M, dropping then significantly from 0.14 to 0.05 when urea concentration increases to 7 M, thus suggesting a relevant decrease of the structural ordering around the tryptophan residues during urea denaturation (Figure 5B).

**Figure 5:**
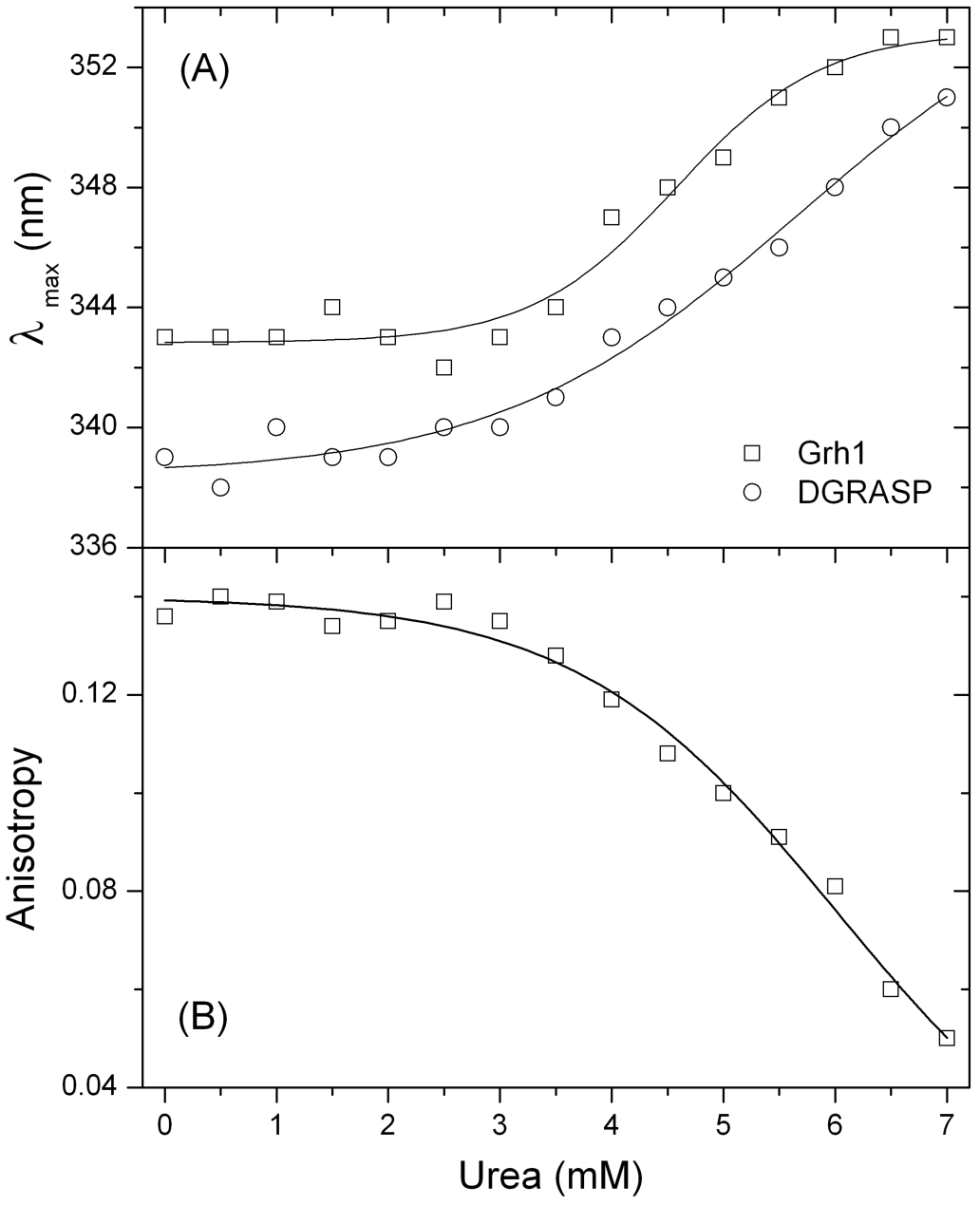
(A) Changes in the emission maxima (λ_max_) and (B) in the steady state anisotropy of Trp fluorescence as a function of the denaturant concentrations. The solid lines are fits using a sigmoidal Boltzmann function.

Since the three tryptophans present in Grh1 are found in the PDZ2, the same fluorescence experiments performed with DGRASP give similar results. However, in this case, the λ_max_ is at 339 nm, a slightly lower value than for Grh1, indicating that the tryptophan residues are less exposed to the solvent as compared to the whole protein. The presence of the SPR domain in the full-length protein seems to induce higher exposure of the inner regions of the PDZ2 domain, suggesting that they do not form two completely separated unities and they may, somehow, interact with each other.

Our observations of the urea-induced unfolding of Grh1 and DGRASP show weak cooperative transitions monitored by CD and somewhat more cooperative unfolding when looking at Trp fluorescence. This apparently disagreement can be explained by the origin of the chromophore under investigation in each method. Far-UV CD measures the optical activity originated from the peptide bonds, whereas fluorescence detects the emission of light generated by specific residues in the protein structure (in our case, Trp residues). We can thus see that CD is reporting unfolding of the overall protein structure, while Trp fluorescence is telling the same story from a more localized point of view. The differences in cooperativity seen from those methods indicate the coexistence of disordered and ordered regions both in Grh1 and in DGRASP, which is in agreement with our CD deconvolution and disorder prediction results (see above). The features observed so far, including low protein compaction but still significant amount of ordered secondary structure and low cooperativity during the unfolding transition, are characteristic of molten globule structures, a behavior already observed for a Grh1 homologue (Mendes et al, 2016). Interestingly, the SPR domain does not seem to be determinant for this, which is an issue still to be addressed in further details. Because the GRASP domain is the most conserved region within the GRASP family (Kinseth et al., 2007), we can strongly suggest that members of this family might all be molten globule-like proteins.

#### 3.2.2. Effects of organic solvents

Based on the results shown in the previous sections, we conclude that Grh1 behaves as a molten globule like protein in solution and presents features attributable to proteins containing multiple intrinsically disordered regions. It has been shown that GRASP from *C. neoformans* (CnGRASP) experiences multiple disorder-to-order transitions upon changes in the dielectric constant of the medium or dehydration (Mendes et al., 2018). GRASP are peripherally associated to membranes, so it is expected that disturbances in the physicochemical parameters induced by biological membranes may have some influence on its structure. A unique disturb induced by the biological membrane is the change in the dielectric constant (ε) nearby its surface (Bychkova et al., 2014; Cherepanov et al., 2003). Typically, a dielectric gradient is observed at the membrane/water interface, which can be modeled by an exponentially increasing function from ε = 2-4 at the first water layer up to 78 at approximately 5-6 nm from the interface (Cherepanov et al., 2003). In order to check whether Grh1 is also affected by those alterations in the medium, we performed CD experiments in the presence of organic solvents as mimetic models for the ε variation.

Figure 6 shows that the shape and intensity of the CD spectrum of Grh1 considerably change in the presence of non-aqueous solvents manifested by the increase in the negative ellipticity around 222 nm. As observed in Table 1, the content of helical structure increases 43% and reaches a maximum in 35% Methanol solution. Grh1 behaves similarly to CnGRASP up to this methanol concentration (Mendes et al, 2018). For further increases in methanol, a distinct pattern is observed: Grh1 gains β–sheet secondary structure and loses disordered regions as methanol increases (Figure 6A). The disordered regions decreased 41% in 45% methanol solution. A similar behavior is observed with high concentrations of ACN that induces β-sheet (23%) and helical (51%) conformations and reduces in 50% the disordered regions (Table 1 and Figure 6B). Hence, the decrease in ε induces the collapsed intrinsically disordered Grh1 to fold in a multiphasic manner, just as described by Uversky (2009) for α-synuclein.

**Figure 6:**
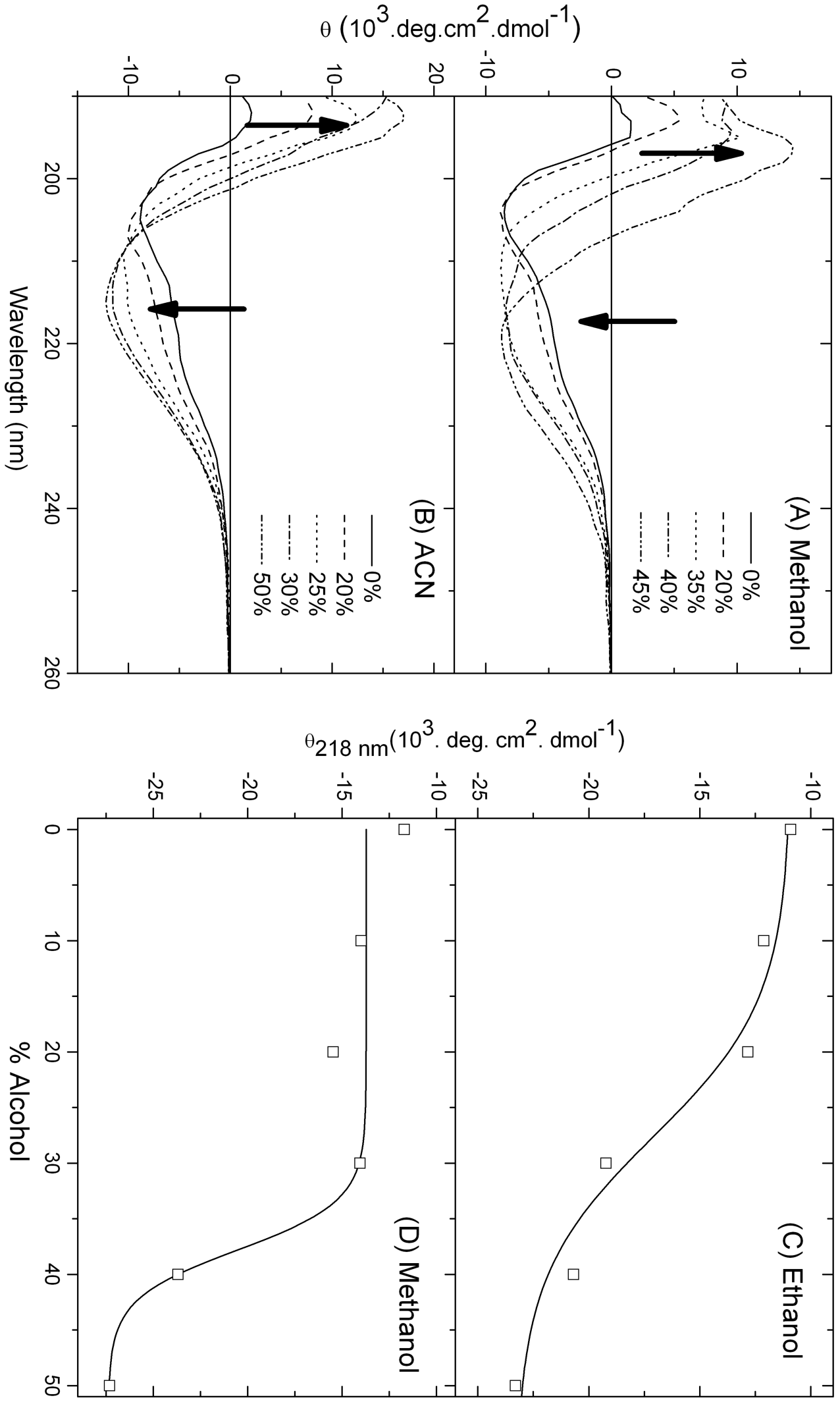
Far-UV CD spectra monitoring the effects of (A) methanol, and (B) ACN on Grh1 structure. Molar ellipticities at 218 nm

**Table 1:**
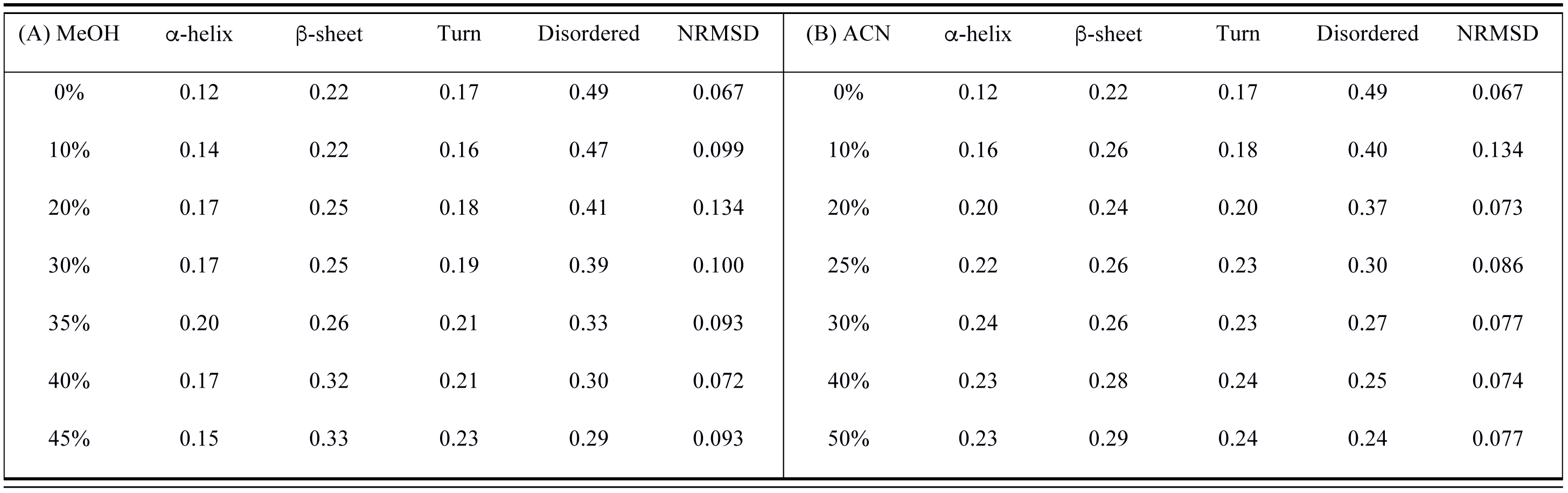
Secondary Structure Content of Grh1 as obtained from deconvolution of the respective CD spectra. The deconvolutions were performed using the Dichroweb software, with the k2d algorithm (Andrade et al., 1993). The data refers to CD spectra measured in increasing concentrations of (A) MeOH and (B) ACN.

| (A) MeOH | α-helix | β-sheet | Turn | Disordered | NRMSD | (B) ACN | α-helix | β-sheet | Turn | Disordered | NRMSD |
| --- | --- | --- | --- | --- | --- | --- | --- | --- | --- | --- | --- |
| 0% | 0.12 | 0.22 | 0.17 | 0.49 | 0.067 | 0% | 0.12 | 0.22 | 0.17 | 0.49 | 0.067 |
| 10% | 0.14 | 0.22 | 0.16 | 0.47 | 0.099 | 10% | 0.16 | 0.26 | 0.18 | 0.40 | 0.134 |
| 20% | 0.17 | 0.25 | 0.18 | 0.41 | 0.134 | 20% | 0.20 | 0.24 | 0.20 | 0.37 | 0.073 |
| 30% | 0.17 | 0.25 | 0.19 | 0.39 | 0.100 | 25% | 0.22 | 0.26 | 0.23 | 0.30 | 0.086 |
| 35% | 0.20 | 0.26 | 0.21 | 0.33 | 0.093 | 30% | 0.24 | 0.26 | 0.23 | 0.27 | 0.077 |
| 40% | 0.17 | 0.32 | 0.21 | 0.30 | 0.072 | 40% | 0.23 | 0.28 | 0.24 | 0.25 | 0.074 |
| 45% | 0.15 | 0.33 | 0.23 | 0.29 | 0.093 | 50% | 0.23 | 0.29 | 0.24 | 0.24 | 0.077 |

In all cases, the CD spectra at the end of the organic solvent variation show a pronounced minimum in the vicinity of 218 nm, typical of folded proteins with β-enriched structures. One can see the transition from α-helical (0% alcohol) to β-rich structures (50% alcohol) in ethanol and methanol (Figure 6C and D) as monitored by changes in the ellipticity at 218 nm. Uversky (2009) described a similar observation when investigating the formation of oligomers of αsynuclein. Considering the fibrillar behavior of α-synuclein depending on the environment (Uversky, 2009) and, being the formation of the β-rich Grh1 irreversible, we hypothesized that the gaining of β-sheet structure could, in fact, be also associated with the formation of fibrils.

#### 3.2.3. Effects of Temperature

Far-UV CD was also used to analyze the thermally induced unfolding of Grh1. Figure 7 represents the far-UV CD spectra of Grh1 measured at different temperatures and shows that the Grh1 spectrum has its shape significantly changed as a function of the temperature. However, the spectra at higher temperatures are not typical of unfolded structures as observed in the thermal unfolding of globular proteins (Kheshinashvili et al., 2016). Instead of reaching a completely unfolded state, Grh1 irreversibly transitioned to a conformation still showing a high content of secondary structure. As the temperature is increased, the minima at 222 nm and at 205 nm become more and less intense, respectively, yielding a mid-point melting temperature (Tm) of 39.1 °C (Figure 7B). Interestingly, it has been previously observed a quite similar result for extended IDPs (Lopes et al., 2013; Uversky, 2009), where it has been proposed that the hydrophobic interactions at higher temperatures are the driving forces for the folding of the polypeptide chain. However, in the previous cases there is a transition from a fully unfolded state to a still unfolded one but with a small increase in helical content, whereas for Grh1 there is a “shape shift” from a folded conformation to a final unknown conformation, which is still rich in ordered secondary structure. Interestingly, the far UV CD spectra progressively undergo a shift to spectra with a minimum at 218 nm, and whose shape and intensity measured at temperatures above 45 °C are close to those recorded in 40% methanol and 50% ACN solutions (Figure 6A and B), showing a β-sheet enriched conformation. Unlike other IDPs, in which temperature effects are reversible (Uversky, 2009), once Grh1 reaches the β-sheet rich conformation, the structure is no longer changed upon cooling. The results in Figure 6 and in Figure 7 suggest that the partial structure disturbances induced by either moderately higher temperature or decrease of ε are sufficient to trigger a transition to an ordered still unknown state of Grh1.

**Figure 7:**
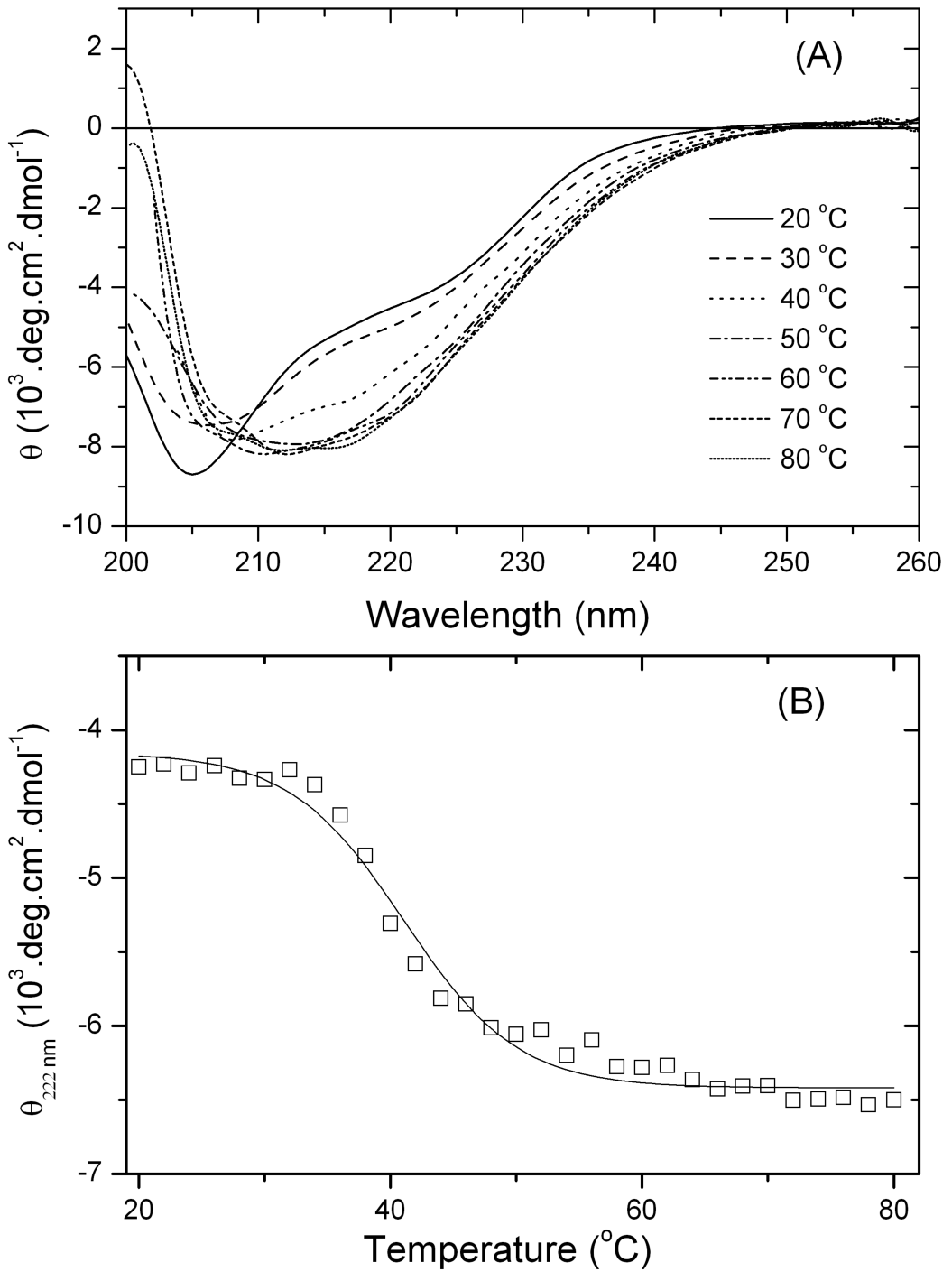
(A) Far-UV CD spectra of Grh1 upon heat-induced unfolding in aqueous solution from 20 to 80 °C. (B) Thermal unfolding monitored by the molar ellipticity values at 222 nm.

Those observations upon changes in temperature and in the presence of organic solvents indicate that, depending on the environment, Grh1 assumes a transient conformation but above a determined threshold, a rigid β-sheet rich conformation is adopted and changes are no longer observed even at high temperatures (up to 80 °C).

### 3.3. Aggregation Prediction and Sequence Analysis

The appearance of considerable b-sheet contributions to the CD spectra of Grh1 either upon heating (Figure 7A) or in the presence of organic solvents (Figure 6) prompted us to investigate whether those b-sheet conformations could be related to aggregation. Similar to the prediction of intrinsic disorder, there are now a number of algorithms to predict the protein regions prone to aggregation. One can have information on aggregation propensities by looking at specific residues that are known to be more common in, for example, amyloid fibrils, such as glutamine and asparagine. The server AGGRESCAN (Conchillo-Solé et al., 2007) evaluates the protein’s primary sequence, classifying the residues in prone or not prone to aggregation. This classification is based not only on the nature of the residue itself, but also on its surroundings (in our case, we chose a five residue window, which means the residue will be evaluated together with the two previous and the two subsequent residues). With that classification, a Hot Spot (HS), a region where 5 or more residues are considered to be prone to aggregation, is created by the server. The longest the region and the aggregating nature of the residues, the higher the HS. The aggregation profile of Grh1 is shown in Figure 8, and we can see a number of short along with three long HS. While the predictor is not exclusive for fibril formation, since other aggregates can exist, it gives a good hint on whether or not a determined region is more likely to form fibrils.

The existence of potential aggregation spots brings the close link between aggregation and intrinsic disorder into play (Wu; Fuxreiter, 2016). Hence, to check for correlations between intrinsic disorder and aggregation in the case of Grh1 we also show in Figure 8 one of the disorder predictions presented in Figure 1. The black solid line represents the threshold for intrinsic disorder. The flexibility gained with a less compact structure can be used to help overcoming energy barriers needed for the formation of the aggregate. Several structural arrangements of disorder and aggregate-like regions in proteins have been proposed (Wu; Fuxreiter, 2016) and in one of them the amyloid core is flanked by intrinsically-disordered regions (IDRs), which could be the geometry adopted by Grh1 as suggested by the intrinsically disorder and aggregation propensities shown in Figure 8.

**Figure 8:**
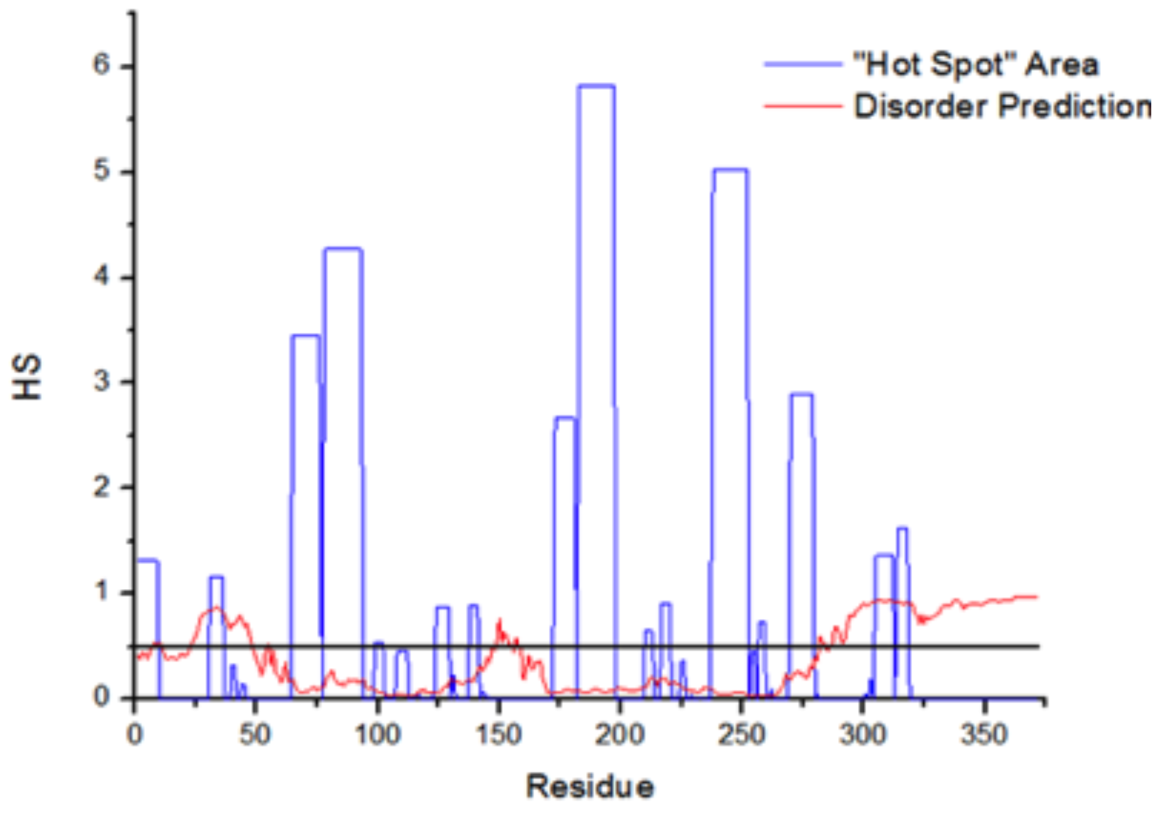
Aggregation prediction for Grh1, done in the AGGRESCAN server, represented by “Hot Spot” areas in blue. The red lines represent one of the disorder predictions showed in figure 1

As for the final residues in the sequence, those in the SPR domain, it is reasonable not to observe aggregation since prolines are considered to be chain breakers, thus leading the score of a determined window in AGGRESCAN to 0. That means a domain such as the SPR would not aggregate. We can also think of that in terms of the structure of the fibril: accommodate a proline into a β-sheet is very costly in terms of energy (Bolognesi et al., 2010).

### 3.4. Ghr1 forms β-sheet rich amyloid fibers

#### 3.4.1. 8-anilino-1-naphthalenesulfonic acid (ANS) assay

Our bioinformatics analysis strongly suggested that Grh1 contains regions that are prone to aggregation, which, in conjunction with our results on the intrinsically disordered nature of part of Grh1 structure, indicate that Grh1 would be able to form β-sheet rich amyloid fibers. The formation of fibrils is a process that includes the formation of small oligomers that associate due to a destabilization of the native structure, leading to the formation of a number of partially folded intermediates, which possess increased aggregation propensity. This process is often called “monomer activation” (Morris et al., 2009). In their review on the modeling of amyloid fibril formation, Gillam and MacPhee (2013) cover the first moments of amyloid formation, called the lag phase, the mechanisms underlying the growth phase, where the formation of the proto-fibrils happens, until their assembly in amyloid fibrils, on the plateau phase. If we look at the whole process during time, we will have a sigmoid-like behavior much like the one we see in our CD experiments (Figure 6C and D).

To further investigate if Grh1 is really forming fibers depending on the environment conditions, we followed the well established protocols based on the use of the fluorescence of extrinsic dyes (Bolognesi et al, 2010). ANS is a fluorescent dye commonly used in protein folding studies (Hawe et al., 2008). Although it is not specific for amyloid fibrils, the experiment we conducted followed previous studies related to fibril formation. Bolognesi et al. (2010) were able to trace all the phases of fibril formation as a function of increasing concentrations of a fibril trigger. Even more interesting in that report, the authors were able to establish a good relation between ANS fluorescence and the presence of proto-fibrils. Since ANS will bind to accessible hydrophobic cores in the protein, when the monomers assemble into proto-fibrils, there will be new hydrophobic sites created, thus increasing ANS fluorescence intensity. Keeping the stimulus by increasing the trigger concentration, the system is forced into the plateau phase, where the proto-fibrils assemble to form the proper amyloid fibrils. By doing so, the fibrils lose hydrophobic sites previously present, and then the ANS fluorescence decays (Bolognesi et al., 2010). Figure 9 shows how the ANS fluorescence will increase with increasing concentrations of ethanol (that we had seen on the CD experiments to lead to aggregation), until it reaches a maximum in 45% ethanol, and then decreases in 50% ethanol, which is in agreement with our CD data (Figure 6C). Although we cannot see the sigmoid-like time-course formation of the fibrils, we have data that is consistent with previous findings regarding ANS binding to proto-fibrils (inset in Figure 9).

**Figure 9:**
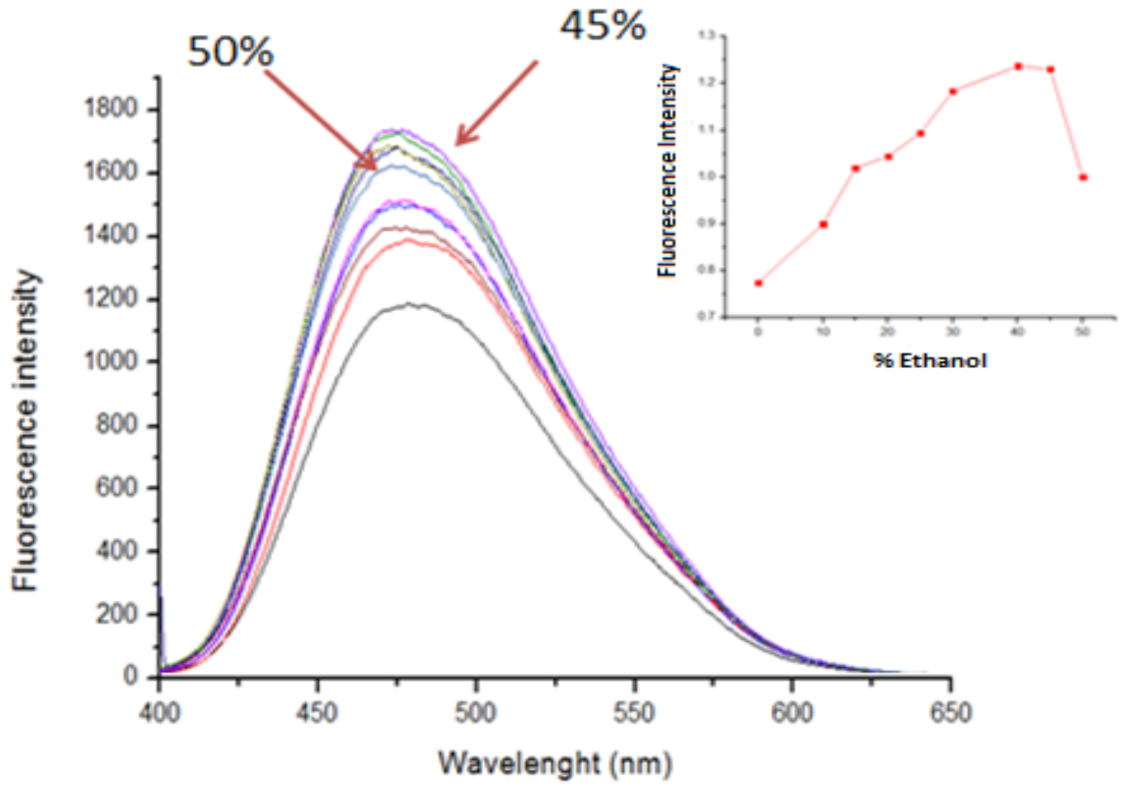
Fluorescence spectra of ANS bound to Grh1 with increasing concentrations of ethanol. The arrows point to the spectra in 45% and in 50% ethanol, emphasizing the reduction in intensity above 45% ethanol.

#### 3.4.2. Thioflavine T (ThT) assay

The ANS assay described in the previous section indicates that Grh1 undergoes a structural transition from monomers to fibrils upon increasing ethanol concentration. To check whether those fibrils present amyloid features, we performed an assay based on the use of the fluorescence of Thioflavine T (ThT) in the presence of Grh1. ThT is a fluorescent dye used in the detection and characterization of amyloid fibrils *in situ* (Nilsson, 2004). It works as a fluorescent rotor that binds into β-sheet cavities (Groenning et al., 2007). When in solution the fluorescence is weak due to ThT freedom of rotation. When bound to fibrils, there is less torsional relaxation, leading to an expressive increase in fluorescence (Nilsson, 2004). Although ThT can bind to amorphous aggregates and other structures with minor affinity, it is considered specific for amyloid fibrils (Nilsson, 2004). In Figure 10A, we see weak fluorescence when ThT is in solution with Grh1 in its native form. However, when ThT is in solution with Grh1 previously submitted to the conditions we have seen to induce aggregation (we tested for temperature, methanol, ethanol and acetonitrile) there is at least a 10-fold increase in the fluorescence intensity. We tested for the same conditions with our construction without the SPR domain (Figure 10B) and we observe the same increase in ThT fluorescence. Although the comparison between the ThT data for Grh1 and the GRASP domain cannot provide any insight into the route of fibril formation, it is nevertheless another proof that the SPR domain is not needed for the fibrillation to occur. Furthermore, we can see that different conditions led to different intensities in ThT fluorescence. For both Grh1 and DGRASP, ACN showed to induce the largest change, while heating led to a large change in DGRASP, but a not so pronounced one for Grh1, in which the effects of ethanol and methanol were markedly more pronounced. That can be the result of either the preparation of the samples not being exactly equal, or it can be related to the pathways and the configuration that each condition induces.

**Figure 10:**
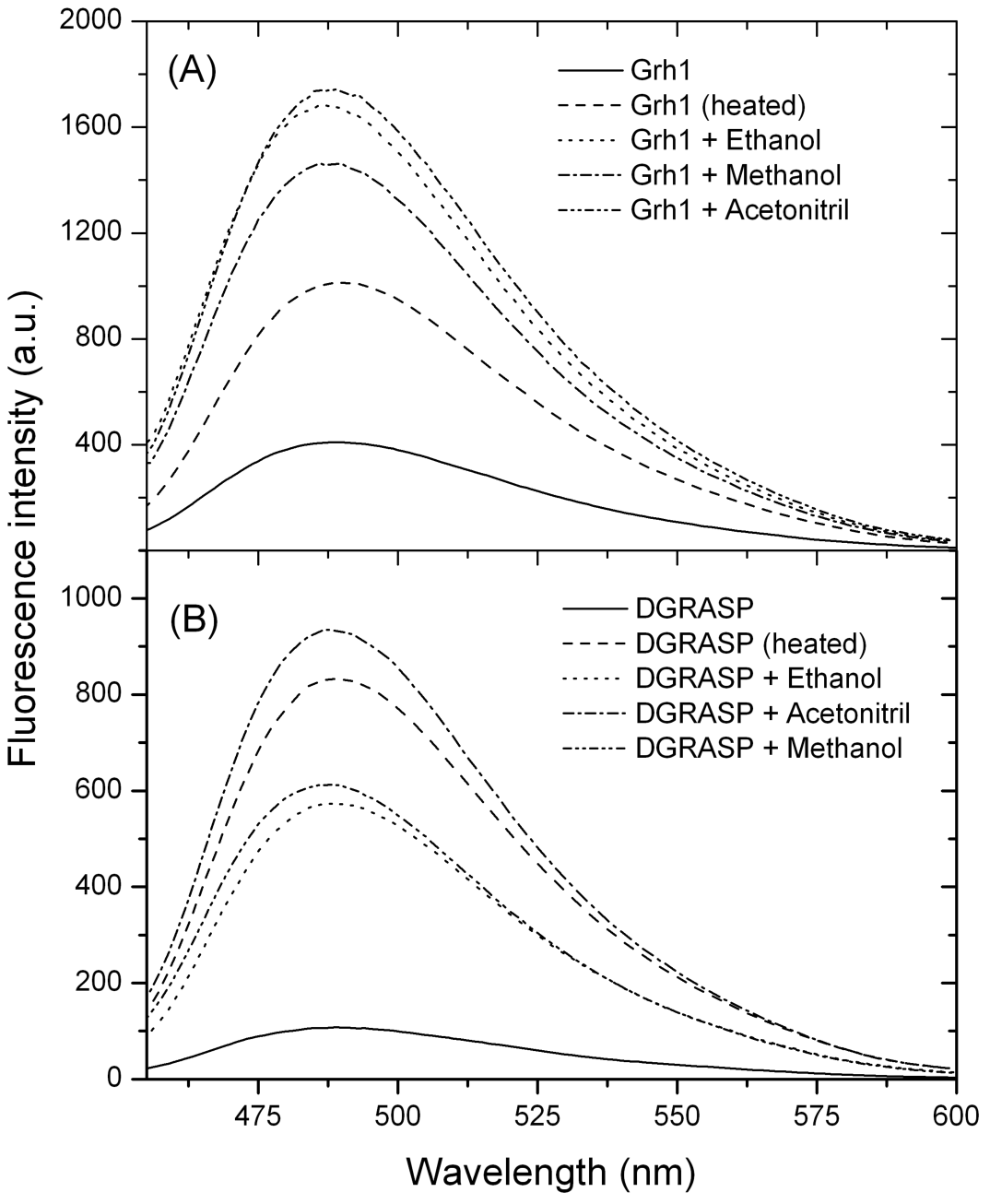
Fluorescence spectrum for ThT bound to (A) Grh1 and (B) GRASP domain only, in different conditions.

#### 3.4.3 Congo Red (CR) assay

CR is another widely used dye to probe amyloid structures (Nilsson, 2004). The exact mechanism of binding is still unknown, but there are some models for it, such as ionic interactions between the sulfonate group of CR and basic residues in the aggregate (Klunk et al., 1999). It is possible to use the birefringence of the amyloid fibrils with CR to prove their existence but, since several participants are inherently birefringent (such as buffer salts), the technique is quite subjective and requires a known amyloid structure as control (Nilsson, 2004). For that reason, we chose another approach based on a spectrophotometric assay. For this experiment we used pre-heated Grh1 to 50 °C to assure fibrillation. In Figure 11 we can see that the absorbance of CR between 400 and 700 nm increases linearly with the increase in pre-heated Grh1. We tested for the native form of Grh1, but there is no change in CR absorbance, showing that the binding only takes place with the protein in its fibrillar (pre-heated) form. Despite the fact that CR binding is one of the most accepted evidences of amyloid formation, it is now known that amyloid fibrils of different compositions may bind to CR through different mechanisms, which can change CR response (Nilsson, 2004). The most pronounced change in absorbance intensity around 540 nm (in this experiment, the largest difference was in 533 nm) is believed to be characteristic of amyloid fibrils.

**Figure 11:**
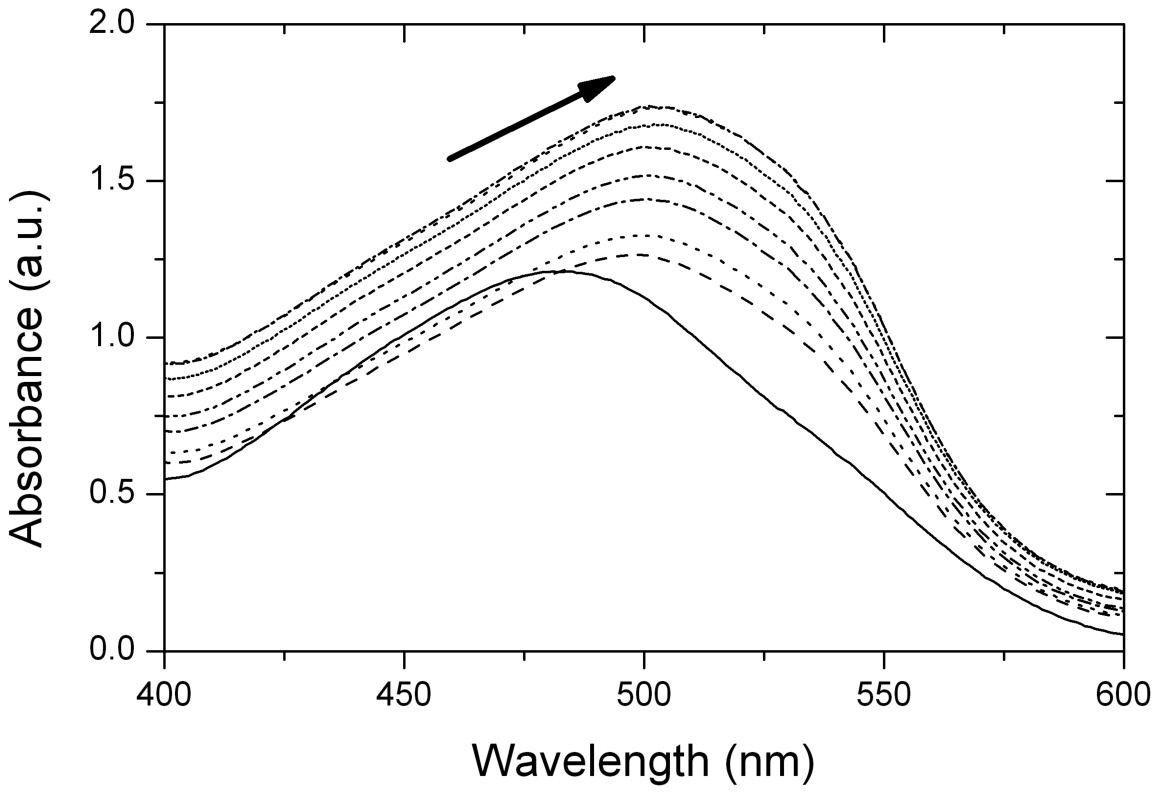
Absorbance spectrum for Congo Red free and bound to pre-heated Grh1.

#### 3.4.4. Transmission Electron Microscopy

The final experiment to check for fibril formation was Transmission Electron Microscopy, which allowed us to actually see the fibrils. Figure 12 shows one of the images obtained from Negative Staining experiments. In this, we are able to see fibrils along with some minor structures, possibly oligomers, which did not have time to overcome the growth phase and become mature fibrils. As expected for amyloid fibrils, our structures are unbranched and present constriction sites but, at that resolution, we were unable to obtain their atomic-resolution structures. Statistical analysis using the program Image J (Schneider et al., 2012) showed the fibrils have a mean size of 55 nm, and are 10 nm wide, with a constriction site of 4 nm. As previously described for other amyloid forming IDPs (Wu and Fuxreiter, 2016), it is common to find IDRs flanking amyloid regions, so it is possible that the SPR domain, although not needed for the fibrillation process to occur, does influence the structure as a whole.

**Figure 12:**
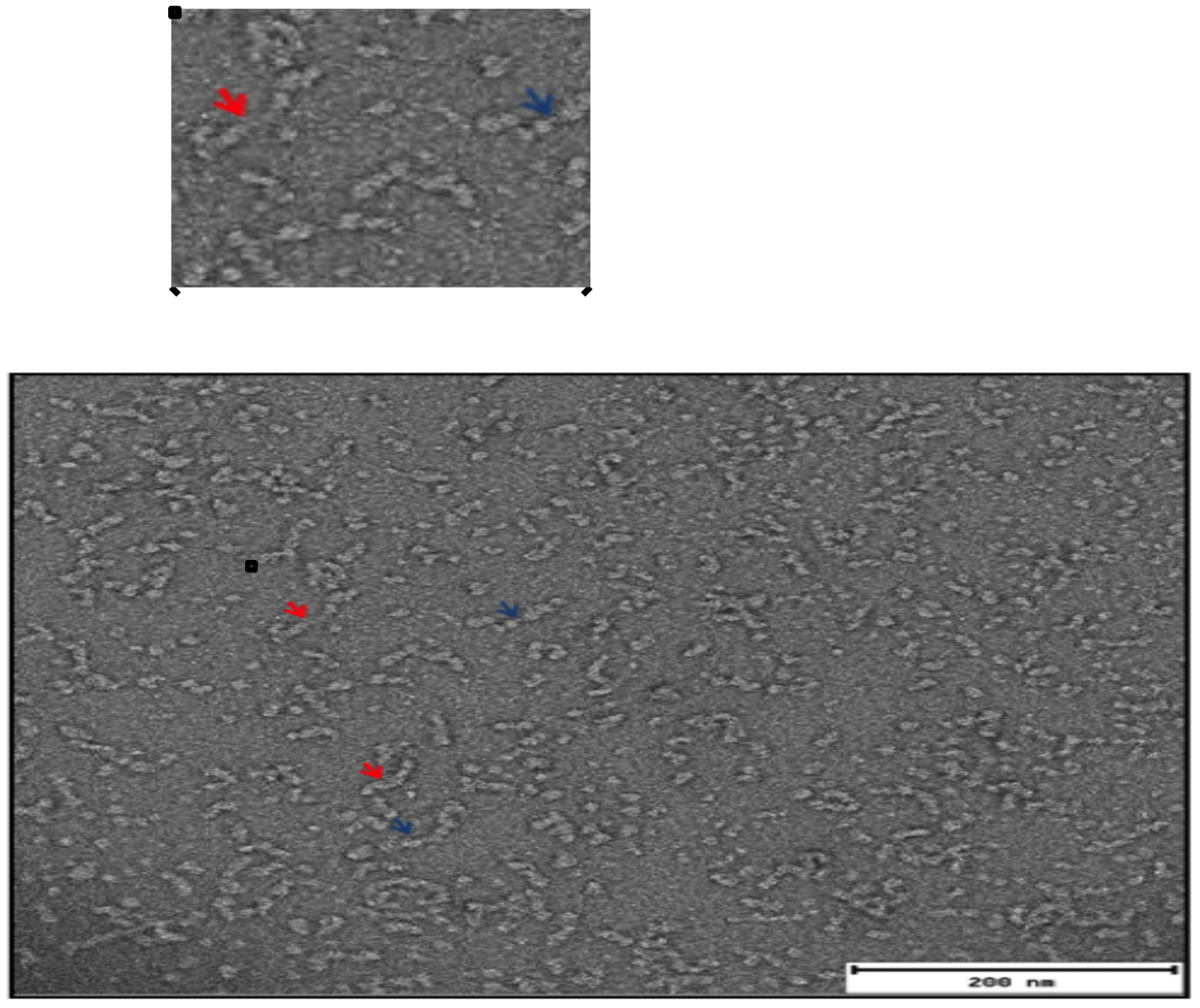
TEM image of Grh1 heated to 45 °C. The red arrows indicate individual fibrils, while the blue arrows indicate the constriction sites

Based on what we have seen on the TEM images, and on what we already know about the Grh1 homolog GRASP65 (Feng et al., 2013), we propose here a model of aggregation. Given that the SPR domain is most likely not part of the fibril itself, we kept it out of our model for simplicity. Considering our results from CD experiments, tracing the signal at 218 nm (Figure 6C and D), and also what is described in the literature as the pathway of aggregation (Landreh et al., 2016), our model suggests a sigmoid-like behavior for the process. The GRASP domain is represented as the two PDZs in ball-shape of different colors (Figure 13). There is also a connecting element representing the constriction site. It is noteworthy that this element is also part of the GRASP domain, probably a loose region, such as a loop, that projects itself away from the core of the PDZ. Given that Grh1 is a homolog of GRASP65, we suppose here the same size seen in structural studies of this human GRASP (Hu et al., 2015). Our model suggests a first step of dimerization, during the growth phase, where 2 molecules of about 6 nm (considering two round PDZs) would account for the ca. 10 nm width of the fibril. Oligomerization would then continue to bring dimers together through the connecting element until the plateau phase is reached. So we have a proper fibril, composed of about 10 dimers, explaining the mean size of 55 nm found in the TEM experiment.

**Figure 13:**
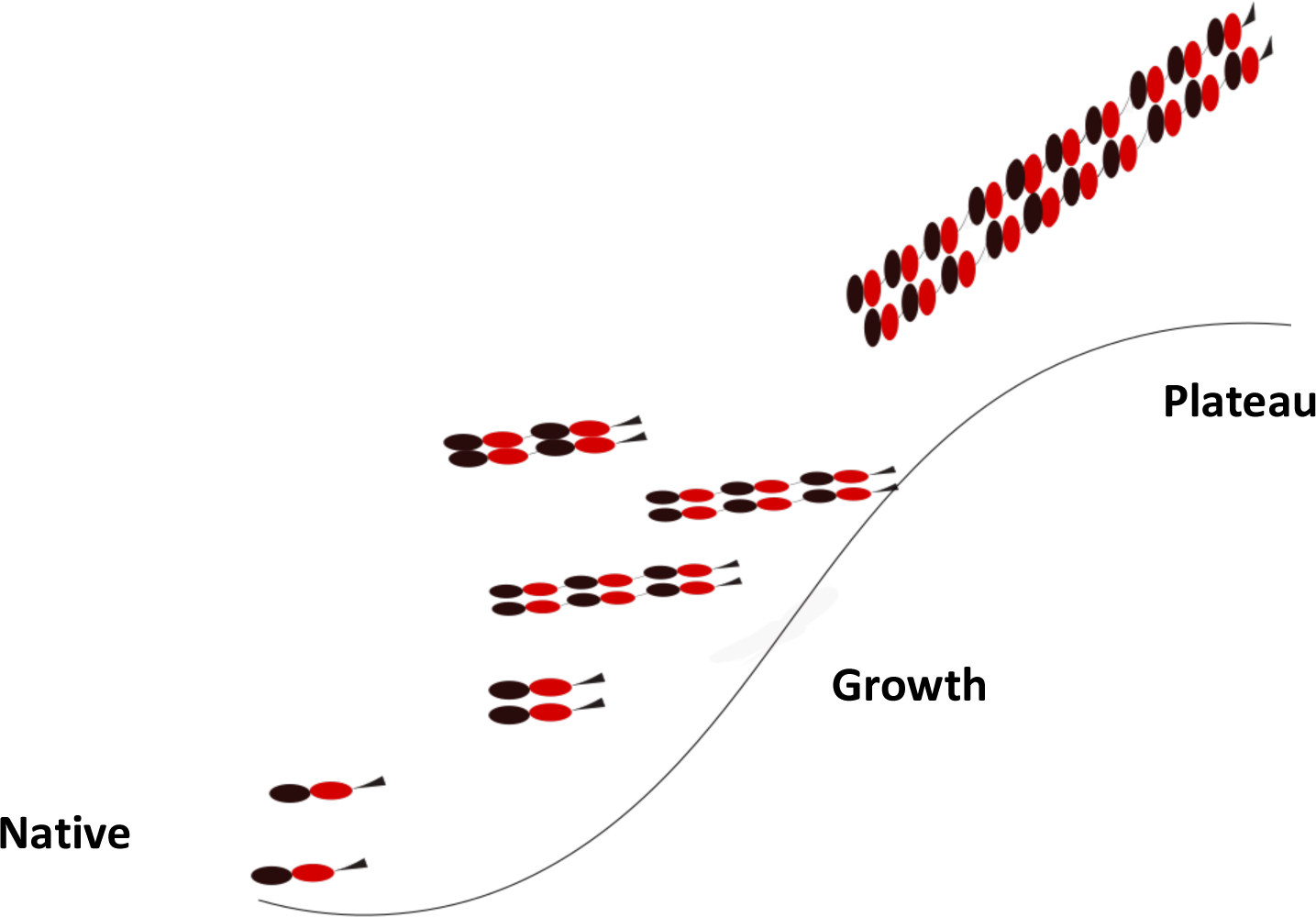
Suggested mode of aggregation: black and red circles represent the GRASP domain of Grh1. In the growth phase, initiated via stimuli such as temperature increase, DGRASP undergoes dimerization. The dimers then assemble into oligomers via the structure denoted here as the constriction site. The process continues until about 10 dimers are joined, reaching then the plateau phase.

## 4. Conclusions

In this manuscript, we have described a biophysical characterization of Grh1 and its GRASP domain that revealed two significant aspects about Grh1, which are most likely linked: the presence of multiple intrinsically disordered regions that confers to Grh1 a molten globule-like feature and the capability of forming amyloid fibrils upon mild denaturing conditions, in an SPR-independent fashion. Grh1 structural dynamic in solution seems to be high but still showing a minimum stable tertiary structure. However, when a destabilizer condition, such as high temperature and/or the membrane surface, is introduced and the structure is slightly disturbed, an irreversible transition associated with amyloid fibril formation is induced. Interestingly, amyloid formation of b2m, a dialysis-related amyloidosis disease resulting from deposition of amyloid aggregates in skeletal tissue, is strongly enhanced in conditions that destabilize its globular structure (Gejyo et al., 1985). Besides, the interaction between α-synuclein and lipids has been also shown to modulate amyloid fibril formation, depending on the relative proportion of the two species (Galvagnion et al. 2015), suggesting that the membrane surface is capable of triggering fibrillation. It has been suggested that partially folded α-synuclein structures induced by increasing temperature is stabilized by self-assembly and that these oligomers may evolve into the fibril nucleus, besides having all the properties expected for a molten globules (Uversky et al., 2001). In general, misfolding intermediates play a key role in defining aberrant protein aggregation and amyloid formation in several different human diseases (Kisilevsky, 200). We observed that the GRASP domain is capable of forming fibbers in a SPR independent way, and since this is the most well conserved region along GRASP family, it is reasonable to expect the same amyloidogenic pattern for all members.

Amyloid fibrils are also found in a diversity of organisms, such as plants and bacteria (Wu; Fuxreiter, 2016), and *Saccharomyces cerevisiae* is not an exception. There are reports of amyloid proteins in yeast, such as the termination factor Nab3, that together with other two proteins forms a complex that is the major termination tool for short, non-coding RNAs (O’rouke; Reines, 2016).

Once thought to be disease-related only, today the idea of functional amyloids is widely accepted. Bacteria and even humans can use the properties of some fibrils to perform functions in the organism (Fowler et al., 2007). Such is the case of Sup35p: yeasts carrying the aggregated form of the protein have selective growth advantage (Fowler et al., 2007). In the case of Grh1, interestingly, only under growth it localizes to ER exit sites and early Golgi membranes, and the yeast stops growing above 37 °C (Dominguez et al., 1982), around the same temperature we determined that Grh1 forms fibrils. Upon stress conditions, like starvation and incubation at the nonpermissive temperature of 37 °C, Grh1 redistributes normally to a large compartment called compartment for unconventional protein secretion (Cruz-Garcia et al., 2014; Duran et al., 2010). Thus, a hypothesis for further investigation is whether the formation of fibrils by Grh1 takes part in membrane fusion events to help generating compartments involved in unconventional secretion (Cruz-Garcia et al., 2014). Because we observed that both temperature and the membrane surface could affect the fibril formation, it is interesting to explore deeply the phenomenon when both perturbations are present together. Experiments to address this are currently being performed.

Understanding the relationships between protein structure and function is one of the fundamental questions in molecular biophysics. We proved that Grh1 is a marginally stable protein and undergoes folding reactions that involve different kinds of ordered forms depending on the environment. The functional diversity reported for Grh1 can then greatly benefit from the possibility of becoming more ordered or folded into stable secondary or tertiary structures and increase the specificity of binding. Furthermore, the irreversible quaternary structure it adopts (the amyloid fibrils) in some conditions might be a strategy of evolution to help survivability in undesired conditions.

## Acknowledgements

The authors are grateful to LNNano (Brazilian Nanotechnology National Laboratory) and the Molecular Biophysics group “Sergio Mascarenhas” for the access to the transmission electron microscopy and circular dichroism equipment, respectively. LFSM and NAF thank Fundação de Amparo à Pesquisa do Estado de São Paulo (FAPESP) for the scholarships (FAPESP Proc. 2012/13309-7, 2016/09676-5 and 2016/23863-2). AJCF thanks FAPESP (Grant nos. 2012/20367-3, 2015/18390-5 and 2015/50366-7) and Conselho Nacional de Desenvolvimento Científico e Tecnológico (CNPq) (Grant no. 308380/2013-4) for the financial support.

---

**Suppl. Material _Figure S1:**
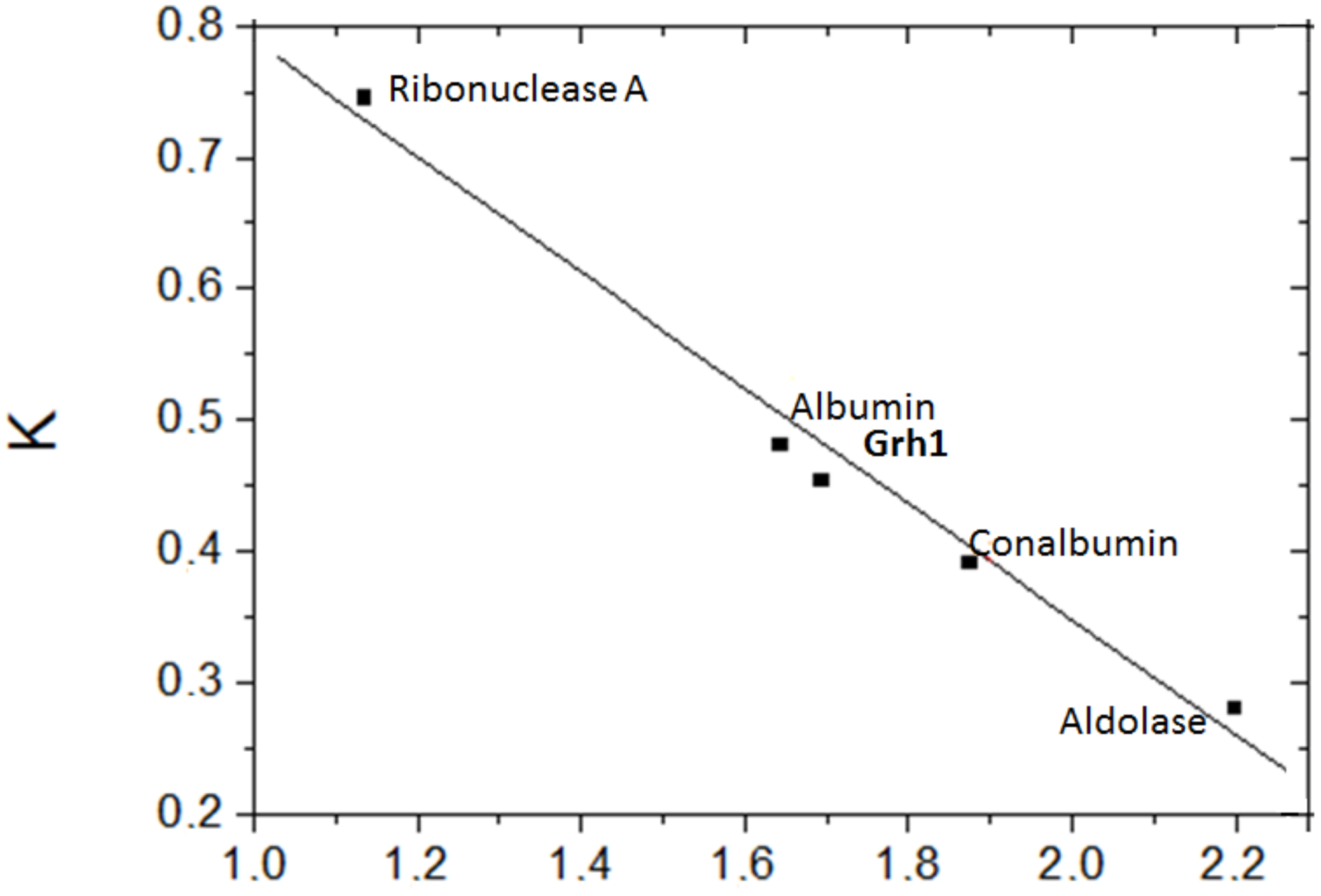
Elution curve. Relationship between the partition coefficient (K) and the logarithm of Molecular Mass

## References

Andrade, MA, Chacón, Merelo JJ, Morán F. Evaluation of secondary structure of proteins from UV circular dichroism using an unsupervised learning neural network. Prot. Eng. 1993. 6, 383–390.

Bachert C, Linstedt AD. Dual anchoring of the GRASP membrane tether promotes trans pairing. J Biol Chem. 2010 May 21;285(21):16294–301.

Barr FA, Puype M, Vandekerckhove J, Warren G. GRASP65, a protein involved in the stacking of Golgi cisternae. Cell. 1997, 91(2):253–262.

Behnia R, Barr FA, Flanagan JJ, Barlowe C, Munro S. The yeast orthologue of GRASP65 forms a complex with a coiled-coil protein that contributes to ER to Golgi traffic. J Cell Biol. 2007 Jan 29;176(3):255–61.

Bolognesi B, Kumita JR, Barros TP, Esbjorner EK, Luheshi LM, Crowther DC, Wilson MR, Dobson CM, Favrin G, Yerbury JJ. ANS binding reveals common features of cytotoxic amyloid species. ACS Chem Biol. 2010 5(8):735–40.

Bychkova VE, Basova LV, Balobanov VA. How membrane surface affects protein structure. Biochemistry (Mosc). 2014 79(13):1483–514.

Chakrabortee S, Tripathi R, Watson M, Schierle GS, Kurniawan DP, Kaminski CF, Wise MJ, Tunnacliffe A. Intrinsically disordered proteins as molecular shields. Mol Biosyst. 2012 8(1):210–9.

Cherepanov DA, Feniouk BA, Junge W, Mulkidjanian AY. Low dielectric permittivity of water at the membrane interface: effect on the energy coupling mechanism in biological membranes. Biophys J. 2003 85(2):1307–16.

Conchillo-Solé O, de Groot NS, Avilés FX, Vendrell J, Daura X, Ventura S AGGRESCAN: a server for the prediction and evaluation of “hot spots” of aggregation in polypeptides. BMC Bioinformatics. 2007 8:65.

Cruz-Garcia D, Curwin AJ, Popoff JF, Bruns C, Duran JM, Malhotra V. Remodeling of secretory compartments creates CUPS during nutrient starvation. J Cell Biol. 2014 207(6):695–703.

Dominguez A, Varona RM, Villanueva JR, Sentandreu R. Mutants of Saccharomyces cerevisiae cell division cycle defective in cytokinesis. Biosynthesis of the cell wall and morphology. Antonie Van Leeuwenhoek. 1982 48(2):145–57.

Dunker AK, Lawson JD, Brown CJ, Williams RM, Romero P, Oh JS, Oldfield CJ, Campen AM, Ratliff CM, Hipps KW, Ausio J, Nissen MS, Reeves R, Kang C, Kissinger CR, Bailey RW, Griswold MD, Chiu W, Garner EC, Obradovic Z. Intrinsically disordered protein. J Mol Graph Model. 2001;19(1):26–59.

Dunker AK, Silman I, Uversky VN, Sussman JL. Function and structure of inherently disordered proteins. Curr Opin Struct Biol. 2008 18(6):756–64.

Duran JM, Anjard C, Stefan C, Loomis WF, Malhotra V. Unconventional secretion of Acb1 is mediated by autophagosomes. J Cell Biol. 2010 188(4):527–36.

Feng Y, Yu W, Li X, Lin S, Zhou Y, Hu J, Liu X. Structural insight into Golgi membrane stacking by GRASP65 and GRASP55 proteins. J Biol Chem. 2013 288(39):28418–27.

Finn RD, Bateman A, Clements J, Coggill P, Eberhardt RY, Eddy SR, Heger A, Hetherington K, Holm L, Mistry J, Sonnhammer EL, Tate J, Punta M. Pfam: the protein families database. Nucleic Acids Res. 2014 42:D222–30.

Fowler DM, Koulov AV, Balch WE, Kelly JW. Functional amyloid‐‐from bacteria to humans. Trends Biochem Sci. 2007 32(5):217–24.

Galvagnion C, Buell AK, Meisl G, Michales TC, Vendruscolo M, Knowles TJP, Dobson CM. Lipid vesicles trigger α-synuclein aggregation by stimulating primary nucleation. Nat Chem Biol. 2015, 11, 229–234.

Gejyo F, Yamada T, Odani S, Nakagawa Y, Arakawa M, Kunitomo T, Kataoka H, Suzuki M, Hirasawa Y, Shirahama T. A new form of amyloid protein associated with chronic hemodialysis was identified as beta 2-microglobulin. Biochem. Biophys. Res. Commun. 1985, 129, 701–706.

Gillam JE, MacPhee CE. Modelling amyloid fibril formation kinetics: mechanisms of nucleation and growth. J Phys Condens Matter. 2013 25(37):373101.

Giuliani F, Grieve A, Rabouille C. Unconventional secretion: a stress on GRASP. Current Opinion in Cell Biology 2011, 23:498–504.

Groenning M, Olsen L, van de Weert M, Flink JM, Frokjaer S, Jørgensen FS. Study on the binding of Thioflavin T to beta-sheet-rich and non-beta-sheet cavities. J Struct Biol. 2007 158(3):358–69

Hawe A, Sutter M, Jiskoot W. Extrinsic fluorescent dyes as tools for protein characterization. Pharm Res. 2008 25(7):1487–99.

Heinrich F, Nanda H, Goh HZ, Bachert C, Lösche M, Linstedt AD. Myristoylation restricts orientation of the GRASP domain on membranes and promotes membrane tethering. J Biol Chem. 2014 289(14):9683–91.

Hu F, Shi X, Li B, Huang X, Morelli X, Shi N. Structural basis for the interaction between the Golgi reassembly-stacking protein GRASP65 and the Golgi matrix protein GM130. J Biol Chem. 2015 290(44):26373–82.

Iakoucheva LM, Dunker AK. Order, Disorder and Flexibility: prediction from protein sequence. Structure. 2003 v. 11, n. 11, p. 1316–1317.

Kazakov AS, Markov DI, Gusev NB and Levitsky DI. Thermally induced structural changes of intrinsically disordered small heat shock protein Hsp22. Biophysical Chemistry 2009, pp. 79–85.

Khechinashvili NN, Kabanov AV, Kondratyev MS, Polozov RV. Thermodynamics of globular proteins. J Biomol Struct Dyn. 2017 14:1–10.

Kisilevsky, R. Amyloids: tombstones or triggers? Nat. Med. 2000, 6, 633–634.

Klunk WE, Jacob RF, Mason RP. Quantifying amyloid by congo red spectral shift assay. Methods Enzymol. 1999 309:285–305.

Kondylis V, Spoorendonk KM, Rabouille C. dGRASP localization and function in the early exocytic pathway in Drosophila S2 cells. Mol Biol Cell. 2005 16(9):4061–72.

Landreh M, Sawaya MR, Hipp MS, Eisenberg DS, Wüthrich K, Hartl FU. The formation, function and regulation of amyloids: insights from structural biology. J Intern Med. 2016 280(2):164–76.

Levi SK, Bhattacharyya D, Strack RL, Austin JR 2nd, Glick BS. The yeast GRASP Grh1 colocalizes with COPII and is dispensable for organizing the secretory pathway. Traffic. 2010 11(9):1168–79.

Mendes LF, Garcia AF, Kumagai PS, de Morais FR, Melo FA, Kmetzsch L, Vainstein MH, Rodrigues ML, Costa-Filho AJ. New structural insights into Golgi Reassembly and Stacking Protein (GRASP) in solution. Sci Rep. 2016 6:29976.

Mendes LF, Basso LGM, Kumagai OS, Fonseca-Maldonado R, Costa-Filho AJ. Disorder-to-order transitions in the molten globule-like Golgi reassembly and stacking protein. Biochimica et Biophysica Acta (BBA) - General Subjects. 2009. https://doi.org/10.1016/j.bbagen.2018.01.009

Morris AM, Watzky MA, Finke RG. Protein aggregation kinetics, mechanism, and curve-fitting: a review of the literature. Biochim Biophys Acta. 2009 1794(3):375–97.

Nilsson MR. Techniques to study amyloid fibril formation in vitro. Methods. 2004 34(1):151–60.

O’Rourke TW, Reines D. Determinants of Amyloid Formation for the Yeast Termination Factor Nab3. PLoS One. 2016 11(3):e0150865.

Permyakov SE, Bakunts AG, Denesyuk AI, Knyazeva EL, Uversky VN, Permyakov EA. Apo-parvalbumin as an intrinsically disordered protein. Proteins. 2008 72(3):822–36.

Perticaroli S, Nickels JD, Ehlers G, Mamontov E, Sokolov AP. Dynamics and rigidity in an intrinsically disordered protein, β-casein. J Phys Chem B. 2014 118(26):7317–26.

Rambourg A, Clermont Y, Hermo L. Three-dimensional structure of the Golgi apparatus. Methods Cell Biol. 1981 23:155–66.

Rambourg A, Clermont Y. Three-dimensional electron microscopy: structure of the Golgi apparatus. Eur J Cell Biol. 1990 51(2):189–200.

Schneider C.A, Rasband W S, Eliceiri KW. NIH Image to ImageJ: 25 years of image analysis. Nature methods 2012, 9(7): 671–675

Shorter J, Watson R, Giannakou ME, Clarke M, Warren G, Barr FA. GRASP55, a second mammalian GRASP protein involved in the stacking of Golgi cisternae in a cell-free system. EMBO J. 1999 18(18):4949–60.

Sreerama N, Woody RW. Estimation of protein secondary structure from circular dichroism spectra: comparison of CONTIN, SELCON, and CDSSTR methods with an expanded reference set. Anal Biochem. 2000 287(2):252–60.

van Stokkum IH, Spoelder HJ, Bloemendal M, van Grondelle R, Groen FC. Estimation of protein secondary structure and error analysis from circular dichroism spectra. Anal Biochem. 1990 191(1):110–8

Theillet FX, Kalmar L, Tompa P, Han K, Selenko P, Dunker AK, Daughdrill GW, Uversky VN. The alphabet of intrinsic disorder, Intrinsically Disordered Proteins. 2013 1:1, e24360.

Truschel ST, Sengupta D, Foote A, Heroux A, Macbeth MR, Linstedt AD. Structure of the membrane-tethering GRASP domain reveals a unique PDZ ligand interaction that mediates Golgi biogenesis. J Biol Chem. 2011 286(23):20125–9.

Uversky VN. Intrinsically disordered proteins and their environment: effects of strong denaturants, temperature, pH, counter ions, membranes, binding partners, osmolytes, and macromolecular crowding. Protein J. 2009 28(7-8):305–25.

Uversky VN, Lee HJ, Li J, Fink AL, Lee SJ. Stabilization of partially folded conformation during alpha-synuclein oligomerization in both purified and cytosolic preparations. J Biol Chem. 2001 276(47):43495–8.

Vinke FP, Grieve AG, Rabouille C. The multiple facets of the Golgi reassembly stacking proteins. Biochem J. 2011 Feb 1;433(3):423–33. doi: 10.1042/BJ20101540. Review. Erratum in: Biochem J. 2011 434(3):575.

Wang Y, Satoh A, Warren G. Mapping the functional domains of the Golgi stacking factor GRASP65. J Biol Chem. 2005 280(6):4921–8.

Whitmore L, Wallace BA. Protein secondary structure analyses from circular dichroism spectroscopy: methods and reference databases. Biopolymers. 2008 89(5):392–400.

Woody RW. Circular dichroism spectrum of peptides in the poly(Pro)II conformation. J Am Chem Soc. 2009 131(23):8234–45.

Wright TA, Stewart JM, Page RC, Konkolewicz D. Extraction of Thermodynamic Parameters of Protein Unfolding Using Parallelized Differential Scanning Fluorimetry. J Phys Chem Lett. 2017 8(3):553–558.

Wu H, Fuxreiter M. The Structure and Dynamics of Higher-Order Assemblies: Amyloids, Signalosomes, and Granules. Cell. 2016 165(5):1055–1066.

Xue B, Dunbrack LR, Williams RW, Dunker AK. PONDR-FIT: A meta-predictor of intrinsically disordered amino acids. Biochim Biophys Acta. 2010 1804(4):996–1010.

